# Niche partitioning and coexistence of sympatric macaques in a fragmented habitat of the Upper Brahmaputra Valley, northeastern India^#^

**DOI:** 10.1101/2022.12.24.521880

**Authors:** Narayan Sharma, Anindya Sinha

## Abstract

How closely related species coexist, especially under conditions of resource limitation, remains an intriguing problem in ecology. Needing to share space and resources, such species are expected to have evolved a variety of mechanisms to reduce competition for these resources. We examined niche partitioning among three sympatric congeners: the northern pig-tailed macaque *Macaca leonina*, rhesus *Macaca mulatta*, and stump-tailed macaque *Macaca arctoides* in Hollongapar Gibbon Sanctuary, a ∼21-km^2^ lowland tropical fragment in the upper Brahmaputra valley of northeastern India. Over 14 months, March 2008 to January 2010, we followed two groups per species, collecting 6,471, 11,062, and 16,723 instantaneous scans on the northern pig-tailed, rhesus, and stump-tailed macaques respectively. We estimated home ranges and their overlap, using minimum convex polygons and 95% kernel density estimates, and compared vertical stratum use, employing the Kruskal-Wallis test. The species’ niche breadth and overlap, using Levins’ and Pianka’s indices respectively, and behavioural activity allocation across six observation time blocks were modelled, using beta-binomial mixed models. Rhesus macaques primarily used forest edges and matrix habitats, their ranges largely separate from those of stump-tailed macaques whereas pig-tailed macaque ranges nested completely, though asymmetrically, within the larger stump-tailed macaque ranges. Vertically, pig-tailed macaques used the highest strata and rhesus macaques the intermediate while the stump-tailed macaques were usually near the ground stratum. Dietary niche breadth was broadest for rhesus macaques and significantly narrower for pig-tailed and stump-tailed macaques, with dietary overlap being highest between rhesus and pig-tailed macaques and lowest between pig-tailed and stump-tailed macaques. Finally, temporal behavioural activity allocation differed significantly between the three species for all five observed behaviours across the day. Considering the three axes of spatial use, including horizontal home range and vertical strata, diet composition, and temporal behavioural activity patterns, therefore, our overall finding that pairs of macaque species that overlap on one of these axes diverge on the others provides evidence for fine-scale partitioning of space and diets between the species, possibly facilitating the coexistence of three congeneric macaque species in this small, highly threatened lowland rainforest fragment.

## Introduction

Understanding how ecologically similar species coexist within the same habitat has remained a central question in community ecology (Elton, 1946; Tokeshi, 1999). The competitive exclusion principle predicts that two species with identical ecological requirements cannot coexist indefinitely within the same habitat (Gause, 1934). Recent theoretical work, however, shows such a pairwise rationale may not always scale to whole communities, wherein indirect interactive mechanisms can permit coexistence even among species that would otherwise exclude one another in isolation (Levine et al., 2017; Miller & Max, 2025). The theory of limiting similarity, on the other hand, proposes that coexisting species must differ sufficiently in their ecological requirements to reduce interspecific competition and persist together within the same community (MacArthur & Levins, 1967). Consequently, ecologically similar or closely related species are expected to partition resources along one or more niche dimensions to facilitate their coexistence.

The niche concept (Hutchinson, 1957), which encompasses the resources and environmental conditions, necessary for the survival and reproduction of a species, remains central to our understanding of multispecies coexistence. Niche partitioning is, therefore, widely regarded as a key mechanism promoting coexistence, particularly through segregation along major niche dimensions, such as habitat use and diet (Schoener, 1974, Coelho et al., 2026). This is especially relevant for closely related or congeneric species because their similarities, arising from shared evolutionary histories, often result in overlaps in morphology, physiology, behaviour or specific ecological requirements, thereby increasing the possibility of interspecific competition (Weber & Strauss, 2016, Coelho et al., 2026). Such partitioning need not, however, necessarily depend on fixed, evolved differences between species, as recent theoretical work has shown that behavioural flexibility in resource use alone can generate sufficient divergence in realised niches for competitors to coexist (Rueffler & Lehmann, 2024).

Ecological niche partitioning has been documented across a wide range of taxa (MacArthur, 1958; Schoener, 1974; Emmons, 1980; Bagchi et al., 2003; Dymit et al., 2025; Azofeifa-Romero, 2026), including nonhuman primates (Terborgh, 1984; Cords, 1986; Ganzhorn, 1989; Schreier et al., 2009). Sympatric primates may partition resources through differential use of habitats or microhabitats (Ganzhorn, 1989; Rodman, 1991; Rakotondranary & Ganzhorn, 2011; Zhou et al., 2014; Justa et al., 2019; He et al., 2024, Howell et al., 2026), vertical strata (Ungar, 1996; Singh et al., 2000; Sushma & Singh, 2006; Singh & Roy, 2011; Feeroz, 2012), dietary resources (Terborgh, 1984; Ganzhorn, 1988; Sushma & Singh, 2006; Ruslin et al., 2019), and behavioural strategies that minimise interactions with heterospecific individuals (Schreier et al., 2009; Justa et al., 2019). Habitat and diet characteristics are considered amongst the most important niche dimensions, influencing coexistence among sympatric species (Schoener, 1974; Zhou et al., 2014, 2026).

The genus *Macaca* is the most widely distributed nonhuman primate genus, spanning habitats from High Atlas Mountains of Morocco in the west to the island of Honshu in Japan in the east, and from mean sea level to over 4,700-m elevations (Fooden, 1982). The genus is also well known for the adaptive radiations displayed by its species, with several sites across South and Southeast Asia supporting two or more sympatric macaque species, thus making the genus an ideal model to study coexistence amongst closely related primates. In this connection, several studies have thus documented ecological niche partitioning between rhesus macaques *Macaca mulatta* and Assamese macaques *M. assamensis* in the Western Himalaya of northern India (Justa et al., 2019), lion-tailed macaques *M. silenus* and bonnet macaques *M. radiata* in the Western Ghats mountains of southwestern India (Singh & Roy, 2011), and between northern pig-tailed macaques *M. leonina* and rhesus macaques in northeastern Bangladesh (Feeroz, 2012) and in southwest China (Sun et al., 2020). However, nonhuman primate assemblages, comprising three sympatric congeneric species, have been considerably less studied (but see Zhou et al., 2026) and the extent to which niche partitioning along spatial, dietary, and temporal dimensions promotes their coexistence remains poorly understood.

Hollongapar Gibbon Sanctuary, situated in Jorhat district of Assam state in the Upper Brahmaputra Valley of northeastern India, is one of the few sites in the world where three congeneric macaque species occur together in the same forest fragment. The sanctuary encompasses 20.98 km^2^ of tropical semi-evergreen and evergreen lowland rainforest, now isolated within a matrix of tea gardens and human settlements, following extensive forest clearance during the late nineteenth and early twentieth centuries CE (Sharma et al., 2012a). Despite its small and fragmented nature, Hollongapar harbours exceptional primate diversity: seven species have been recorded historically, of which six persist today, including three *Macaca* species: *Macaca mulatta, Macaca leonina*, and *Macaca arctoides*, alongside the western hoolock gibbon *Hoolock hoolock*, capped langur *Trachypithecus pileatus*, and the Bengal slow loris *Nycticebus bengalensis* (Sharma et al., 2012b). This extraordinary assemblage of sympatric primates in a fragment of area less than 21 km² makes the sanctuary a unique site for investigating the mechanisms that allow congeneric species to coexist under spatially constrained and resource-limited conditions. This is particularly important as tropical forests, home to over 90% of primate species, are becoming increasingly fragmented globally and now largely exist in small sized patches, with the persistence of many large-bodied mammals dependent on whether and how they can coexist with other sympatric species (Marsh & Chapman, 2013; Haddad et al., 2015; McKinney, 2025).

Here, we investigate ecological niche partitioning among rhesus macaques *Macaca mulatta*, northern pig-tailed macaques *Macaca leonina*, and stump-tailed macaques *Macaca arctoides* in the Hollongapar Gibbon Sanctuary along three primary axes: spatial use, including horizontal home range and vertical strata, diet composition, and temporal behavioural activity patterns. We specifically ask: (1) Do the three macaque species differ significantly in their use of forest space, including both horizontal range and vertical strata? (2) Do they partition their diets by food type, plant species, or plant parts? (3) Do they differ in how they distribute their behavioural activities across the day? By characterising niche partitioning in this three-species assemblage within a human-modified, fragmented landscape, our study contributes to a broader understanding of the conditions under which nonhuman primate diversity can be maintained in forest fragments, with direct implications for the conservation management of Hollongapar and similar degraded rainforest patches across northeastern India.

## Methods

### Study Area

The Hollongapar Gibbon Sanctuary, located in the Jorhat district of Assam state, northeastern India, was declared a Reserved Forest in 1881 and a Wildlife Sanctuary in 1997. The sanctuary was once an integral part of the extensive foothill forests of the Patkai range of the further eastern state of Nagaland. The Hollongapar forest began to fragment and lost its connectivity with the adjacent Patkai foothill forests after the establishment of extensive tea gardens during the colonial period of 1880–1920. The original area of the sanctuary was 20.98 km^2^; approximately 3 km^2^ has, however, since been leased out and now lacks any kind of tree cover. As a result, the effective size of the sanctuary is currently *c*.17.98 km^2^, a quarter of which is highly degraded and also devoid of any canopy. Moreover, Hollongapar is now intersected by a railway line that has effectively fragmented its core area (Fig. 1).

**Fig. 1.**
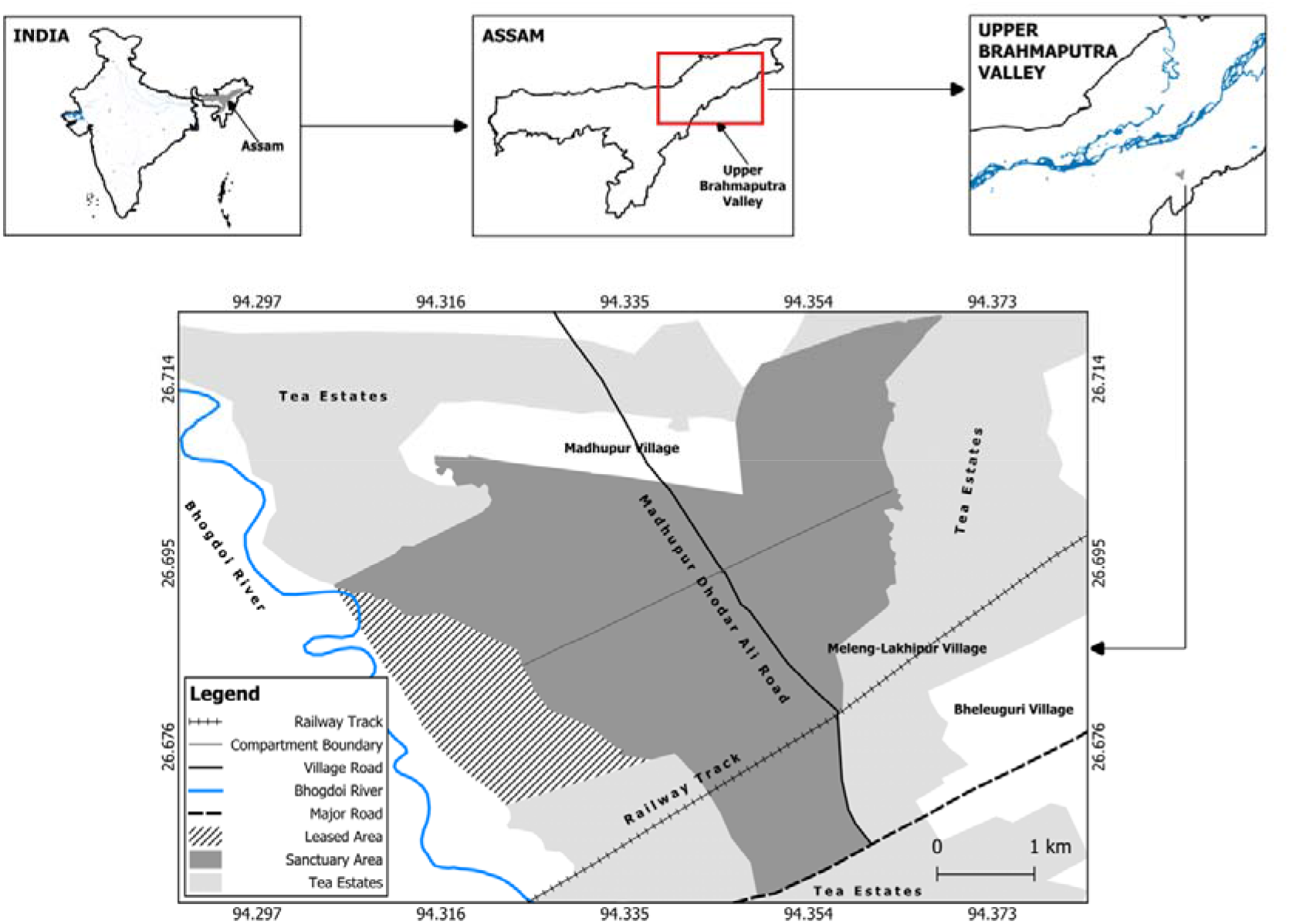
Map of the study area, Hollongapar Gibbon Sanctuary, in northeastern India, March 2008, showing the sanctuary area, leased area, internal demarcations, and the surrounding areas, including tea estates and villages.

The original vegetation type of the sanctuary, which falls under the North East India Biogeographic Zone (9) and NE Brahmaputra Valley Biogeographic Province (9A) (Khatri, 2009), was Assam Plains Alluvial Semi-Evergreen Forests, with sporadic wet evergreen forest patches (Champion & Seth, 1968). The sanctuary, whose elevation ranges from 110 to 120 m ASL, has had a long management history and almost two-thirds of its area is now covered by artificially regenerated forests (Duarah & Saikia, 2003; Khatri, 2009). Today, Hollongapar fragment lies in an extensive mosaic of tea gardens, agricultural fields, and human settlements. The marginalised communities of these settlements largely harvest non-timber forest produce and occasionally fell trees within the sanctuary (Sharma et al., 2020).

Based on the rainfall received, the study period was divided into two seasons: the dry season, from October to March, with a mean monthly rainfall of <100 mm and the wet season, between April and September, with an average monthly rainfall of >100 mm. The mean maximum and minimum temperatures in the sanctuary during the study period were 28°C and 19°C respectively.

### Study Species and Groups

There were originally seven species of nonhuman primates in Hollongapar (Srivastava et al., 2001), of which the Assamese macaque became locally extinct since 2005 (Sharma et al., 2012b). Of the remaining species, the Bengal slow loris is a nocturnal prosimian while the remaining five species—the rhesus macaque, northern pig-tailed macaque, stump-tailed macaque, capped langur, and the western hoolock gibbon—are diurnal. Amongst the macaques, the rhesus macaque is the most ecologically versatile and widely distributed, occurring across all eight northeastern states of India and on both banks of the river Brahmaputra (Choudhury, 2001). The northern pig-tailed macaque occurs in northeastern India, Bangladesh, Myanmar, Thailand, Laos, Vietnam, and China, but in northeastern India, it is restricted to the south bank of the Brahmaputra (Choudhury, 2001). Similarly, the stump-tailed macaque is distributed across South and Southeast Asia, with the river Brahmaputra marking the westernmost limit of its distribution (Choudhury, 2001).

The populations of the five diurnal species in Hollongapar have not only stabilised, but also increased in abundance, with Sharma et al. (2012a) reporting very high densities of all the species in the sanctuary. Hoolock gibbons have had the highest group density (1.4 groups/km^2^) while stump-tailed macaques have occurred at the highest individual density (12.96 individuals/km^2^, Sharma et al., 2012a); the largest group ever of stump-tailed macaques across its distribution range, with 194 individuals, has also been seen in Hollongapar (Sharma et al., 2021).

Two groups of each of the three macaque species were selected for our study, conducted from March 2008 to January 2010. However, the actual behavioural observations were initiated only in November 2008 and continued till January 2010. The intervening period between March 2008 and November 2008 was utilised to select the study groups and mutually habituate to one another. The group size and age-sex classification of the study groups are presented in Table I.

**Table I.** Age-sex composition of the study groups of the three macaque species at the Hollongapar Gibbon Sanctuary, March 2008.

|  | **Group Code** | **Group Size** | **Adult Male** | **Adult Female** | **Subadult** | **Juvenile** | **Infant** | **Unidentified** |
|---|---|---|---|---|---|---|---|---|
| **Pig-tailed macaque** | PTM1 | 29 | 4 | 8 | 3 | 3 | 6 | 5 |
|  | PTM2 | 20 | 4 | 4 | 5 | 5 | 2 | 0 |
| **Rhesus macaque** | RM1 | 20 | 5 | 6 | 4 | 1 | 4 | 0 |
|  | RM2 | 54 | 8 | 10 | 11 | 12 | 7 | 6 |
| **Stump-tailed macaque** | STM1 | 102 | 20 | 25 | 17 | 22 | 18 | 0 |
|  | STM2 | 131 | 18 | 30 | 23 | 30 | 30 | 0 |

### Data Collection

#### Bhehavioural Observations

Instantaneous scan sampling (Altmann, 1974) was conducted on all the visible adult males and females of each group in six 2-h time blocks, beginning from dawn, at ∼05:00 h until dusk, at ∼17:00 h, on each observation day, when the individuals were most active. Each scan lasted for a mean (± SD) of 8.9 (± 2.1) min, with a median of 10 min and range 1–20 min. Scan duration depended largely on the size of the study group, its location, and the prevailing environmental conditions. Only the visible members of each group were scanned, as the groups were mostly large and the individuals often widely spread out, sometimes even up to a radius of 500 m in the case of pig-tailed macaques. It was, therefore, virtually impossible to detect all the group individuals, given the short time available for each scan, which was set to a maximum of 20 min in order to maximise the number of individuals observed in each scan. Successive scans began at a median interval of 20 min (mean ± SD = 20.3 ± 10.2 min, n = 2,806 intervals), with 90% of the inter-scan intervals falling between 15 and 20 min, leaving a median gap of 10 min between two scans. Both scan duration and inter-scan interval were similar across the three study macaque species.

Our scan sampling yielded 2,950 group scans across 144 group-days, including 987 for pig-tailed, 1109 for rhesus and 854 for stump-tailed macaques. The total number of scans on the three species differed among the six 2-h time blocks, with 57 between 05:00 and 07:00 h, 433 between 07:00 and 09:00 h, 636 between 09:00 and 11:00 h, 680 between 11:00 and 13:00 h, 693 between 13:00 and 15:00 h, and 451 between 15:00 and 17:00 h. Each of our scans recorded a mean (± SD) of 11.6 (± 7.0, range 1–47) adult individuals while the actual number of such individuals scanned varied with group size; there were thus 6.6 (± 3.6) individuals of pig-tailed, 10.0 (± 3.4) of rhesus, and 19.6 (± 6.4) of stump-tailed macaques in an average scan on the three species.

The following data were collected during each scan: Date and time of each scan, age and sex of the scanned individual – adult male, adult female or adult individual of unidentified sex, behaviour displayed, including Foraging, Feeding, Resting, Moving, Social Interactions, and Other Behaviours, individual height – on a tree, shrub, bamboo or on the ground, tree and shrub height, individual prey item – invertebrates or vertebrates, food plant species and the part/s fed on, and the habit and phenophase of the food plant. We broadly apportioned the various behavioural activities into the following broad categories:

- Foraging: Active search for any plant material or animal –invertebrate or vertebrate – prey species underneath leaves in the canopy or at any other level of a tree, in split bamboo stems, and/or in ground leaf litter
- Feeding: Handling and ingestion of plant and/or animal food item/s
- Resting: Period of inactivity, including sleeping
- Moving: Any locomotory behaviour, resulting in a change of spatial position of an individual
- Social Interactions: Allogrooming and other affiliative behaviour, agonistic behaviour, reproductive behaviour, and play behaviour
- Other Behaviour: Autogrooming, becoming vigilant, startling, vocalising, body shaking, yawning, sneezing, urinating, defecating

The total number of individual scan records for the northern pig-tailed macaque, rhesus macaque, and stump-tailed macaque were 6,471, 11,062, and 16,723 respectively while the total contact hours, when the structured observations were conducted, for the three species were comparably 352, 387, and 336 hours respectively (Table II).

**Table II.** Sampling effort on the three macaque species in Hollongapar Gibbon Sanctuary, November 2008 – January 2010.

| **Species** | **Group Code** | **Group Size** | **Individual Scan** | **Total Contact Hour** |
|---|---|---|---|---|
| **Pig-tailed macaque** | PTM1 | 29 | 4,394 | 254 |
|  | PTM2 | 20 | 2,077 | 98 |
| **Rhesus macaque** | RM1 | 20 | 6,395 | 226 |
|  | RM2 | 54 | 4,667 | 161 |
| **Stump-tailed macaque** | STM1 | 102 | 11,931 | 259 |
|  | STM2 | 131 | 4,792 | 77 |

#### Group Location

The location of each study group was taken at five-min intervals during the observation period by recording the physical coordinates of the approximate centre of the group with the help of a handheld GPS (Garmin Map 60Cx, Taiwan). At the end of each observation day, all the location data were downloaded using the software GPS Trackmaker. As locations were recorded across the full 14-month observation period, the resulting home-range estimates are likely to be representative of each group’s ranging behaviour (Jacobson et al., 2024).

### Data Analysis

#### Home Range

We estimated the home range of each study group by two methods: Minimum convex polygons (MCP), constructed with the convex hull tool in QGIS *v. 3.34.6*, and fixed kernel density estimation (KDE), determined using kernel utilisation distributions, generated with a reference bandwidth (*h = href*) in the *adehabitatHR* package *v.0.4.22* (Calenge, 2024) in R, from which we extracted the 95% utilisation contours. For both methods, we calculated pairwise intersections between all 15 pairs of the six study groups, representing intra-and interspecific combinations, using the Intersect tool in QGIS. Home range overlap was calculated and expressed as the percentage of a focal group’s home range that intersected those of the other groups, yielding a directional overlap matrix for MCP- and KDE-derived ranges.

#### Height Use

Vertical stratum use was evaluated from the scan records of individual height (m above ground) and compiled across both all scans and independently from scans, coded as Foraging and as Feeding; scans that recorded individuals on the ground were excluded from the analysis. Species differences were first tested with Kruskal–Wallis tests, followed by Dunn’s post-hoc tests with Bonferroni correction, with rank-based effect sizes estimates reported as epsilon-squared, ε² = *H*/(n – 1), where *H* is the Kruskal–Wallis test statistic and n the total number of observations across all groups combined.

As the scans were nested within only two groups per species, scan-level tests of this kind risk pseudoreplication, a drawback frequently highlighted in nonhuman primate behavioural studies (Waller et al., 2013; Whitehouse et al., 2024). We, therefore, refitted each comparison as a generalised linear mixed model (Tweedie family, log link; *glmmTMB*), with species as a fixed effect and group identity as a random intercept, so that the group scan rather than the individual scan served as the effective unit of replication. The significance of species differences was assessed by likelihood-ratio test against a null model and pairwise contrasts obtained from estimated marginal means (*emmeans*) with Bonferroni correction.

#### Dietary Niche Breadth

The dietary niche breadth of all the three study macaques was measured using Levins’s index (Hurlbert, 1978; Krebs, 1999) thus:

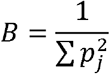

where, *B* = Levins’ index as a measure of niche breadth, and *p_j_* = Fraction of items in the diet that are of food category *j*

This index of niche breadth was standardised to express it on a scale of 0 to 1, as proposed by Hurlbert (1978):

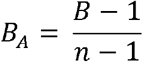

where, *B =* Levins’ index of niche breadth, *B_A_ =* Levins’ standardised index of niche breadth, and n = Number of possible resource states

The food resources, considered in this analysis, included food plant species, plant parts, and vertebrate or invertebrate animal prey. All animal prey species were treated as a single category. If the macaques were observed to feed on more than one part of the same plant species, however, they were considered distinct food resources.

#### Dietary Niche Overlap

Dietary niche overlap between macaque species pairs was quantified, using Pianka’s Index (Pianka, 1973; Krebs, 1999). This index, which ranges from 0 (no overlap in resource use) to 1 (complete overlap), was calculated as:

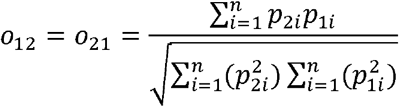

where, O_12_ = Pianka’s index of dietary niche overlap between Species 1 and 2, p_1i_ = Proportion of Resource i in the total resources, used by Species 1, p_2i_ = Proportion of Resource i in the total resources, used by Species 2, and n = Total number of resources

#### Time-Activity Budgets and Daily Behavioural Activity Patterns

Time-activity budgets were calculated as the percentage of scans that the adult individuals of each study group spent in the six behavioural categories, listed above, over the study period.

As individual macaques could not be reliably identified for any of the three study species, the scan records of all observed adults were aggregated by species, group, date, time block (explained below), adult sex, and the observed behavioural category to evaluate the daily or temporal activity patterns of that species; we counted all records in each behavioural category within each sampling unit. As mentioned earlier, we used behavioural scan data from the six 2-h time blocks, extending from 05:00–07:00 h in the morning until 15:00–17:00 h in the afternoon.

We fitted a separate generalised linear mixed model for each of five principal behavioural activities: Foraging, Feeding, Resting, Moving, and Social Interactions, with a two-column response of focal-activity counts versus all other counts; all Other Behaviour contributed to the non-focal category. All models used a beta-binomial error distribution, with a logit link to accommodate overdispersion, fitted in *glmmTMB v. 1.1.8* (Brooks et al., 2017) in R *v. 4.3.3* (R Core Team, 2024) thus:

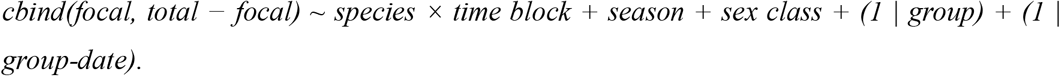

As with vertical stratum use, nested group and group–date intercepts accounted for repeated scans within a day and repeated sampling across days, making the group the effective unit of replication; we tested the group-level variance component against a model omitting it, reporting *p* as a 50:50 chi-squared mixture, given the boundary condition.

Species differences in daily behavioural activity patterns were tested by likelihood-ratio tests, comparing full and no-interaction models, with 10 degrees of freedom and with *p*-values adjusted across the five counted behavioural activities, using the Holm procedure. Where the interaction was significant, we obtained estimated marginal proportions with 95% confidence intervals and Holm-adjusted pairwise species contrasts within each time block, using *emmeans v. 1.10.0* (Lenth, 2024). As the time block of 05:00–07:00 h was sampled relatively inadequately, particularly for pig-tailed macaques, we refit all models after excluding this time block, resulting in eight degrees of freedom. All full and sensitivity models converged, with a convergence code of 0 and positive-definite Hessian. Significance was evaluated at α = 0.05.

## Results

### Home Range

Home range size varied considerably both among and within species (Table III). Following the MCP method, the two pig-tailed macaque groups had the smallest and most comparable ranges—159.38 and 153.58 ha—whereas the rhesus and stump-tailed macaque groups were larger and far more variable: rhesus macaque: 193.66 and 591.05 ha; stump-tailed macaque: 914.03 and 288.76 ha. The largest observed range, that of STM1, was approximately six times the range of either pig-tailed macaque group. The kernel density estimates were similar to the respective MCP values for the pig-tailed macaque groups but diverged significantly for the rhesus macaques, being less than half for the RM1 range (80.99 vs 193.66 ha) but considerably larger for the RM2 range (751.99 vs 591.05 ha).

**Table III.** Home range size and overlap of the six macaque groups in the Hollongapar Gibbon Reserve, estimated using the Minimum Convex Polygon (MCP) and 95% Kernel Density (KD) methods.

| **Species Home Range** |  |  | **Percentage Overlap in Home Ranges** |  |  |  |  |  |
| **Group** | **MCP (ha)** | **KDE (ha)** | **PTM1** | **PTM2** | **RM1** | **RM2** | **STM1** | **STM2** |
|---|---|---|---|---|---|---|---|---|
| **PTM1** | 159.38 | 141.37 | 100<br>(100) | 14.3<br>(16.3) | 6.7<br>(7.8) | 1.9<br>(29.5) | 91.8<br>(96.3) | 0.0<br>(0.0) |
| **PTM2** | 153.58 | 158.95 | 14.8<br>(14.5) | 100<br>(100) | 0.0<br>(0.0) | 0.0<br>(3.9) | 100.0<br>(96.9) | 3.6<br>(17.7) |
| **RM1** | 193.66 | 80.99 | 5.5<br>(13.7) | 0.0<br>(0.0) | 100<br>(100) | 100.0<br>(100.0) | 0.6<br>(11.8) | 0.0<br>(0.0) |
| **RM2** | 591.05 | 751.99 | 0.5<br>(5.5) | 0.0<br>(0.8) | 20.2<br>(10.8) | 100<br>(100) | 0.3<br>(9.0) | 0.0<br>(5.3) |
| **STM1** | 914.03 | 917.54 | 16.0<br>(14.8) | 16.8<br>(16.8) | 0.1<br>(1.0) | 0.2<br>(7.4) | 100<br>(100) | 5.5<br>(9.7) |
| **STM2** | 288.76 | 464.28 | 0.0<br>(0.0) | 1.9<br>(6.1) | 0.0<br>(0.0) | 0.0<br>(8.6) | 17.4<br>(19.2) | 100<br>(100) |

The values provide the percentage of the row group’s home range, estimated using MCP, which fell within the column group’s range. The values within parentheses indicate overlap, estimated using 95% KDE.

The interspecific home range overlaps differed markedly among the species pairs (Fig. 2). The ranges of both pig-tailed macaque groups lay largely within that of STM1, with 91.8% of PTM1’s and 100% of PTM2’s MCP-determined ranges falling within it; the overlap was, however, strongly asymmetric, with the reciprocal values being much lower: 16.0% and 16.8% respectively. The overlap of PTM1 and PTM2 with the second stump-tailed macaque group, STM2 was negligible, being 0.0% and 3.6% respectively. In contrast, the pig-tailed and rhesus macaque groups overlapped little in horizontal space (overlaps of 0.0–6.7% in either direction) while the stump-tailed and rhesus macaque groups were almost entirely separate (overlaps ≤ 0.6%). The KDE method provided relatively higher values for several of these pairs, most notably PTM1 with RM2 (29.5% vs 1.9%) and RM1 with STM1 (11.8% vs 0.6%), indicating peripheral, low-intensity overlaps that the MCP method did not capture.

**Fig. 2.**
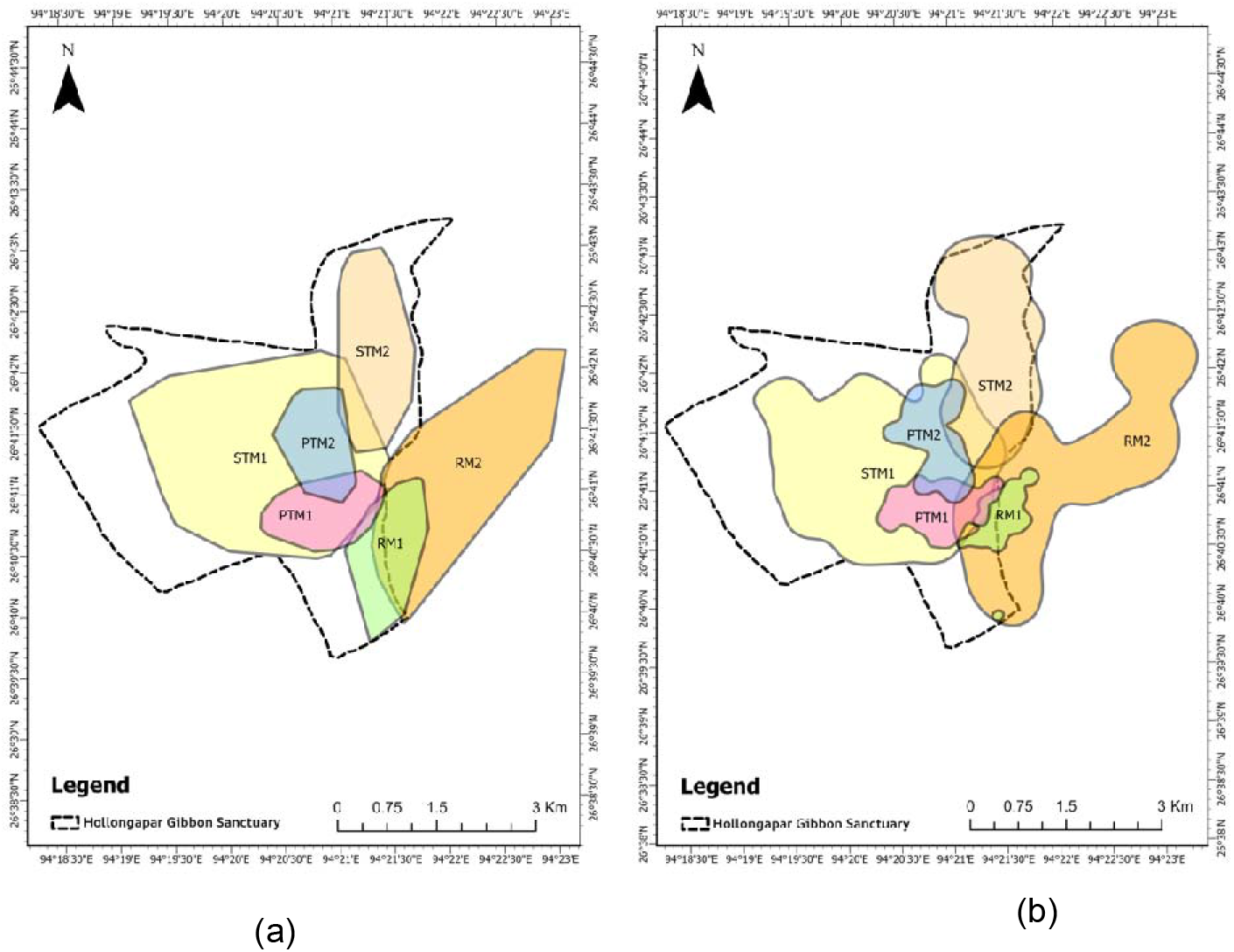
Home ranges of the study groups of northern pig-tailed macaques, stump-tailed macaques, and rhesus macaques in the Hollongapar Gibbon Sanctuary. The home ranges were estimated by both (a) the MCP method and (b) the KDE method.

The intraspecific range overlaps were equally variable (Fig. 2). The entire MCP range of RM1, for example, fell within that of RM2 although the reciprocal value was only 20.2%, reflecting the large range of the latter group. The rhesus macaques were distinctive in mostly occurring on the forest edges and in the surrounding matrix habitat of human settlements and tea plantations. The two pig-tailed macaque groups overlapped moderately and near-symmetrically (14.3% and 14.8%) while there was low overlap between the two stump-tailed macaque groups (5.5% and 17.4%).

### Height Use

Considering overall height use and those used during foraging and feeding, pig-tailed macaques used, on an average, the highest strata, rhesus macaques intermediate strata, and stump-tailed macaques the lowest, near-ground strata (Fig. 3). Kruskal-Wallis tests confirmed significant species differences in all the three datasets (Overall: H = 11264.93, df = 2, *p* < 0.001, ε² = 0.331; Foraging: H = 717.13, df = 2, *p* < 0.001, ε² = 0.741; Feeding: H = 2875.93, df = 2, *p* < 0.001, ε² = 0.275;) while Dunn’s post-hoc tests established all pairwise comparisons significant (all p < 0.001–0.003).

**Fig. 3.**
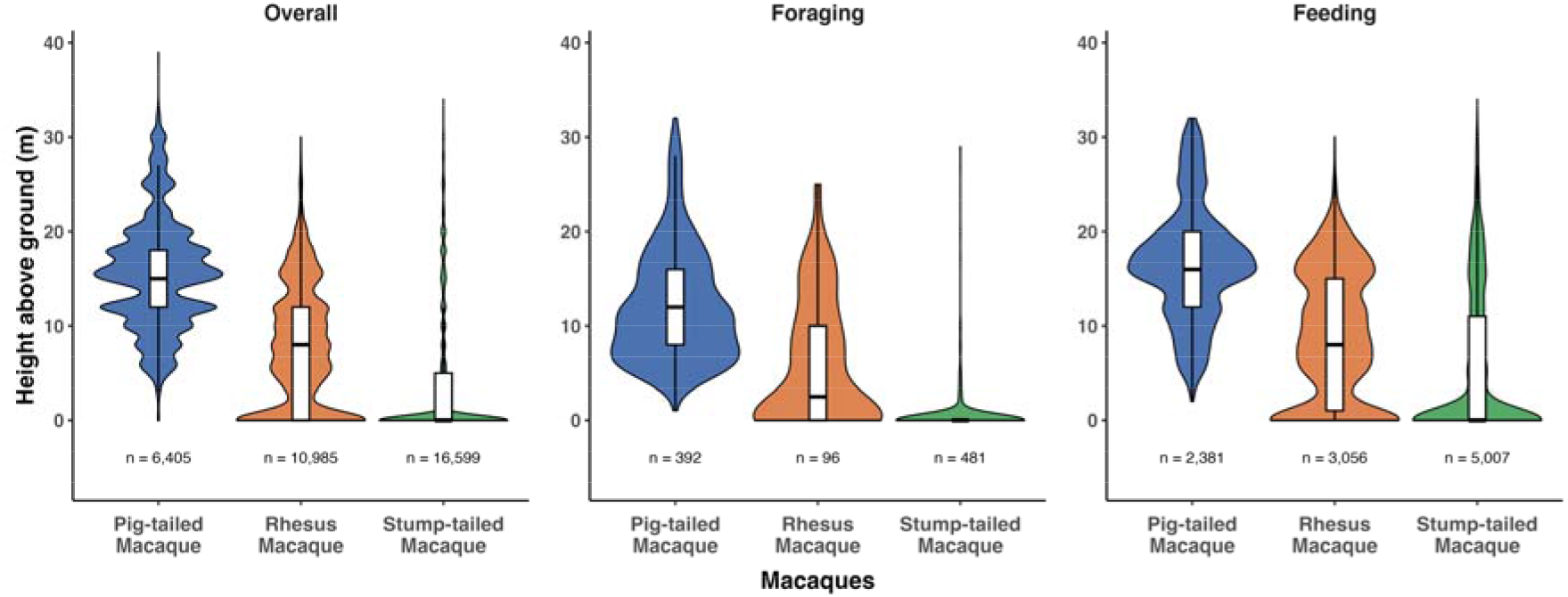
Overall vertical height use and that during Foraging and Feeding (in m above ground) of pig-tailed macaques, rhesus macaques, and stump-tailed macaques in the Hollongapar Gibbon Sanctuary. The violin plots show the full distribution of scan-sampled heights; the overlaid boxplots indicate median and interquartile range. Note the near-zero distribution for stump-tailed macaques during Foraging, reflecting predominantly terrestrial foraging behaviour in this species.

The GLMMs, accounting for group-level non-independence, confirmed a significant species effect in all three datasets (Overall: χ² = 14.58, df = 2, *p* < 0.001; Foraging: χ²(2) = 12.17, df = 2, *p* = 0.002; Feeding: χ² = 17.48, df = 2, *p* < 0.001;), with estimated marginal means closely matching the raw medians (Supplementary Tables S1A and S1B). The pairwise contrasts remained significant for all the species pairs in the overall and feeding datasets as well (Supplementary Table S1C). In the foraging dataset, however, the pig-tailed– rhesus macaques contrast was not significant after group correction (*p* = 0.367) although both species remained clearly distinct from stump-tailed macaques (*p* < 0.001).

### Dietary Niche Breadth and Overlap

We compared the niche breadth and overlap in the dietary profiles of the three macaque species in the dry and wet seasons across the study period. In the dry season, pig-tailed and rhesus macaques had a relatively greater proportion of fruits in their diets (Fig. 4) while the stump-tailed macaque spent a higher proportion of time feeding on root cortices. During the wet season, however, fruits constituted the major part of the diet for all the species. Stump-tailed and rhesus macaques fed significantly on leaves in both the seasons but leaves constituted only a minor part in the diet of pig-tailed macaques across the seasons.

**Fig. 4.**
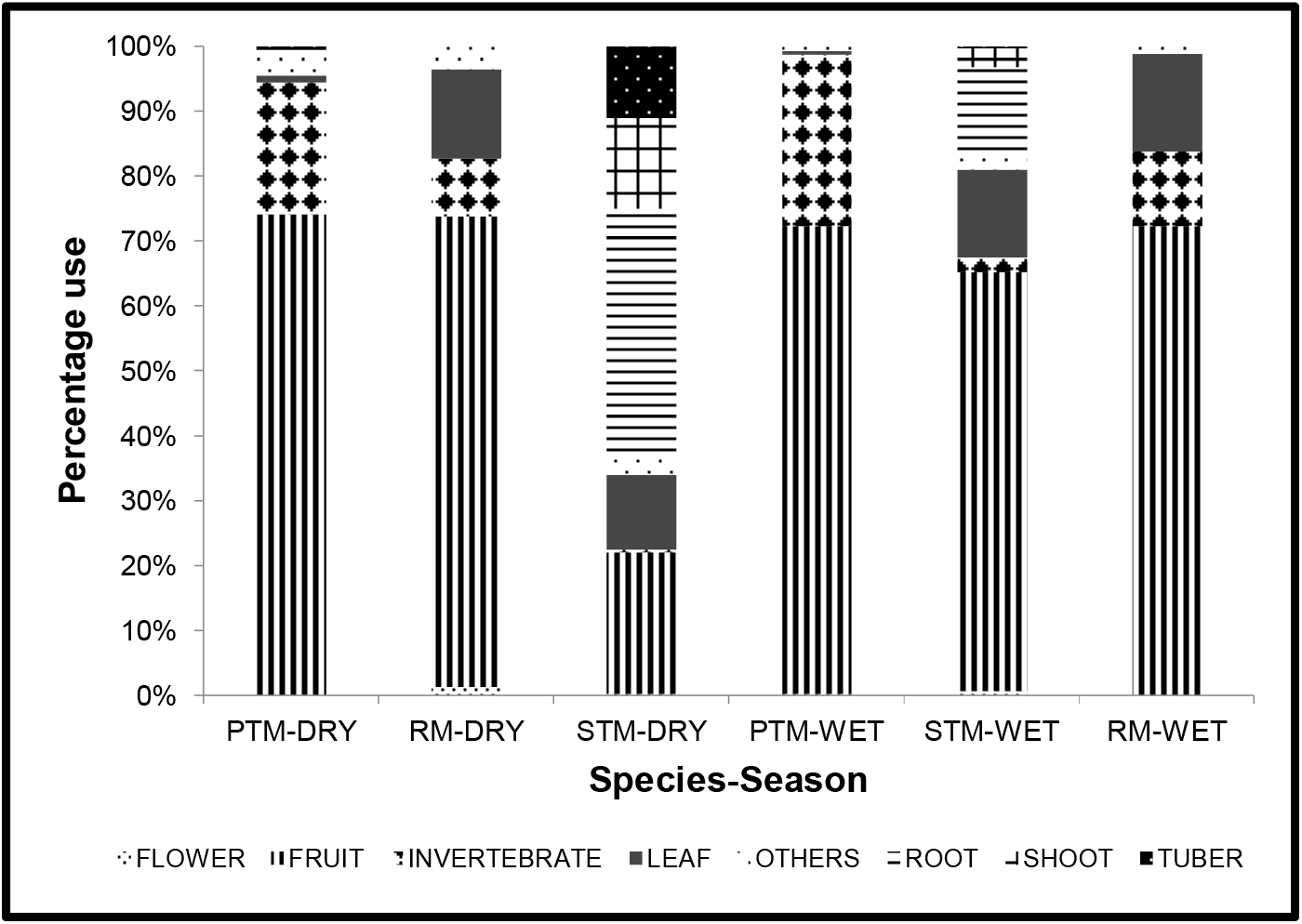
Proportion of different items in the diet of the three macaques in the Hollongapar Gibbon Sanctuary across the dry and wet seasons. PTM: Pig-tailed macaque; RM: Rhesus macaque; STM: Stump-tailed macaque; DRY: Dry season; WET: Wet season.

The standardised dietary niche breadth index value of 0.25 was highest for the rhesus macaque while it was relatively small for both the pig-tailed and the stump-tailed macaque (0.10 and 0.09 respectively; Fig. 5). Moreover, the niche breadth of pig-tailed and rhesus macaques were comparable across the wet and dry seasons but, surprisingly and contrary to what may have been expected, that of the stump-tailed macaque was larger in the wet season than in the dry season.

**Fig. 5.**
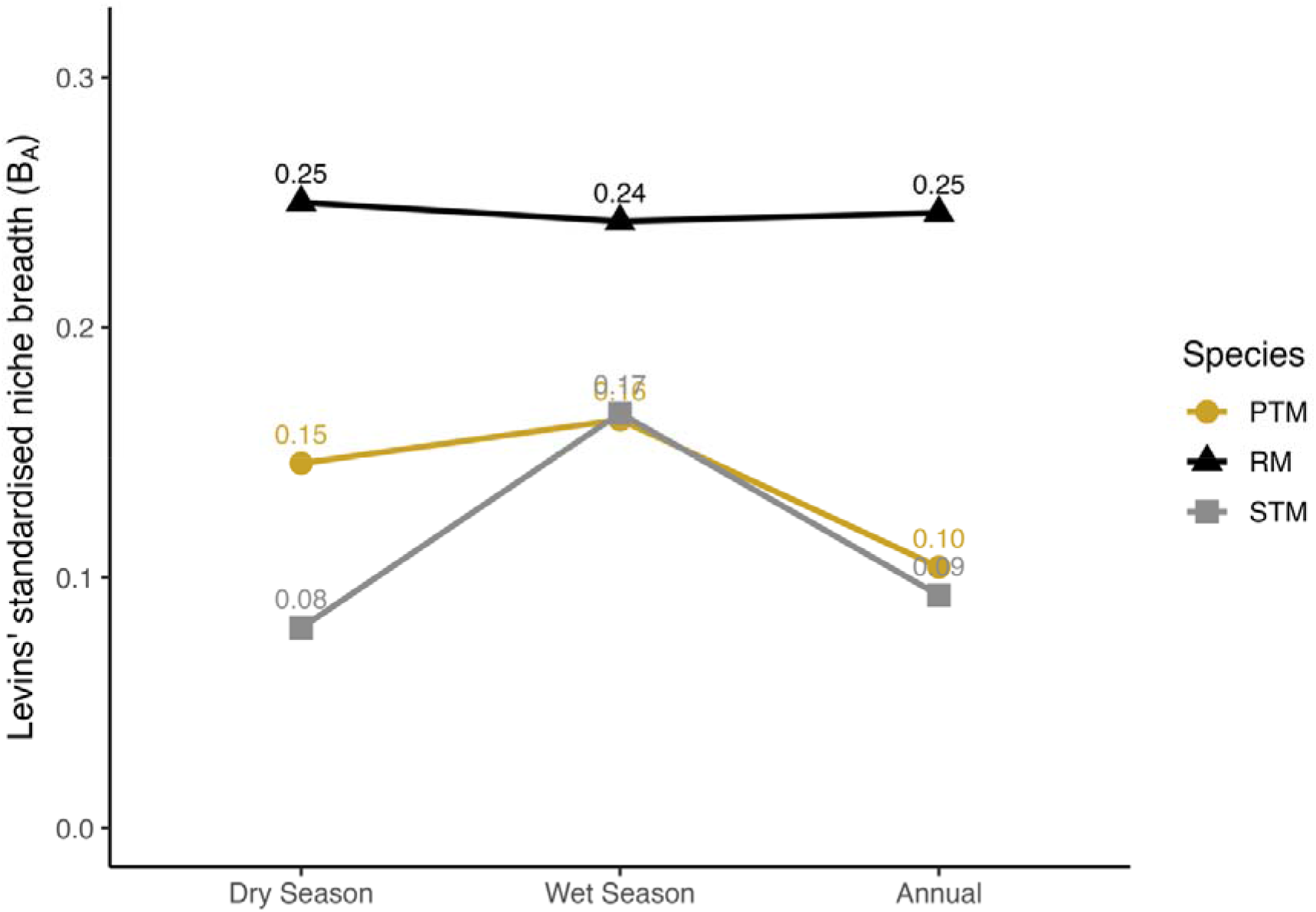
Seasonal and annual dietary niche breadth of the three study macaque species in the Hollongapar Gibbon Sanctuary. PTM: Pig-tailed macaque; RM: Rhesus macaque; STM: Stump-tailed macaque

The pig-tailed macaque and the rhesus macaque had the highest annual overlap in the niche breadth of their diets (0.55), followed by that between the rhesus and stump-tailed macaques (0.18); the diet overlap was least between the pig-tailed and stump-tailed macaques (0.11). There was also a marked seasonal difference in dietary niche breadth overlap. Rhesus macaques had the highest overlap with stump-tailed macaques during the wet season (0.44) but with pig-tailed macaques during the dry season (0.61). Pig-tailed macaques, on the other hand, had very low overlap with stump-tailed macaques, both in the dry and wet seasons (Fig. 6).

**Fig. 6.**
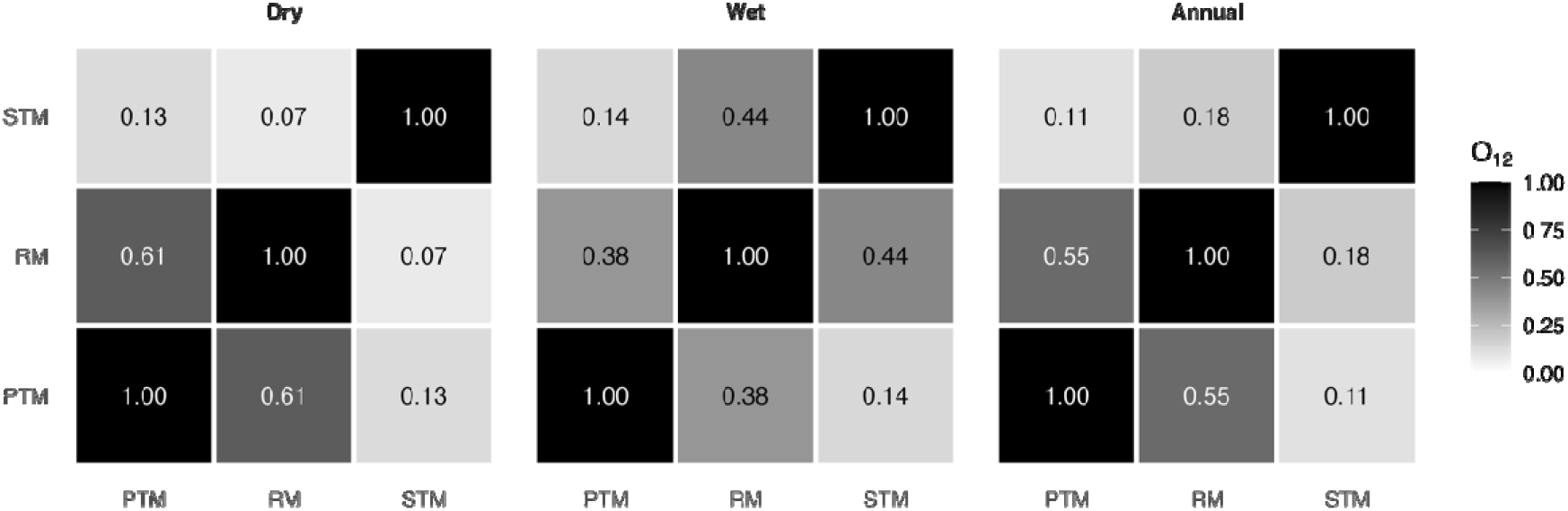
Seasonal and annual Pianka’s niche overlap index (O_12_) values for dietary profiles of the study macaque species pairs in the Hollongapar Gibbon Sanctuary. PTM: Pig-tailed macaque; RM: Rhesus macaque; STM: Stump-tailed macaque

### Time-Activity Budgets and Daily Behavioural Activity Patterns

The time-activity budgets differed amongst the study macaque species groups across the study period (Supplementary Table S2). The daily behavioural activity patterns, represented by the species × time-block interactions were, however, significant for all five behavioural activities: Foraging, Feeding, Resting, Moving, and Social Interactions (Likelihood-ratio test,

Holm-adjusted, *p* ≤ 0.0017 for all activities; Table IV), with Social Interactions showing by far the largest species × time-block interaction effect and Foraging the smallest. These interactions remained significant in a sensitivity analysis, which excluded the sparsely sampled 0500–0700 h period (Table IV), indicating that the detected temporal differences in activity patterns were not driven significantly by observations from the earliest time block.

**Table IV.** Likelihood-ratio tests for the species × time-block interactions in beta-binomial mixed models of daily behavioural activity allocation. The models included group identity and group-date identity as nested random intercepts. The full models included six time blocks (0500–1700; df = 10); the sensitivity models excluded the sparsely sampled 0500–0700 h observation period (five time blocks; df = 8). The *p*-values are Holm-adjusted across the five behaviours within each model set.

| **Behaviour** | **Full Model** |  |  | **Sensitivity Model** |  |  |
|---|---|---|---|---|---|---|
|  | **χ²<sup>\*</sup>** | **df<sup>†</sup>** | **Holm-adjusted *p*<sup>‡</sup>** | **χ²** | **df** | **Holm-adjusted *p*** |
| **Foraging** | 28.25 | 10 | **0.00164** | 27.24 | 8 | **0.000642** |
| **Feeding** | 64.72 | 10 | **9.16 × 10□¹□** | 39.00 | 8 | **9.83 × 10□□** |
| **Resting** | 99.73 | 10 | **2.46 × 10□¹□** | 97.99 | 8 | **4.40 × 10□¹□** |
| **Moving** | 84.85 | 10 | **1.68 × 10□¹³** | 64.01 | 8 | **2.27 × 10□¹□** |
| **Social Interactions** | 195.14 | 10 | **8.31 × 10□³□** | 156.85 | 8 | **3.64 × 10□²□** |

<sup>\*</sup>χ² = Chi-square estimate; <sup>†</sup>df = Degrees of freedom; <sup>‡</sup>*p*-values in bold indicate significance at p < 0.05.

Moreover, group-level variance in behavioural activities was small, relative to variation among group-days, but was greater than zero for Feeding (Group SD = 0.20, p = 0.023) and Social Interactions (Group SD = 0.27, *p* = 0.037), marginal for Foraging (Group SD = 0.24, *p*= 0.099) and negligible for Moving (Group SD = 0.07, *p* = 0.284) and Resting (Group SD = 0.0002, *p* = 0.500; Supplementary Table S3). The species × time-block interactions were nevertheless unaffected by the inclusion of the group-level intercept, changing by less than 2% for four of the five behaviours. Thus, the three macaque species differed not only in their overall time-activity patterns but also in how they distributed their behavioural activities across the day, and these differences were robust to group-level non-independence.

## Discussion

It is truly significant that the three macaque species—the northern pig-tailed, rhesus, and stump-tailed macaques—differed significantly in their use of forest space, in their dietary profiles, and in the timing of their daily behavioural activities in the Hollongapar Gibbon Sanctuary. Horizontal space partitioning was strongest between the rhesus and stump-tailed macaques whereas the pig-tailed and rhesus macaques showed the greatest overlap in diet, even as they diverged in vertical space use and behavioural activity timing over the day. In each of these cases, a pair of macaque species that overlapped substantially on one niche axis proved to differentiate along another, a pattern that, together with distinct daily time-activity budgets across all three species, points to spatial, dietary, and temporal niche partitioning as a plausible mechanism promoting the coexistence of these three congeneric macaques in the study area, one of the important rainforest fragments of the Upper Brahmaputra Valley in northeastern India.

### Spatial Differentiation Among the Three Macaque Species

Horizontal space partitioning, we found, was strongest between the stump-tailed and rhesus macaques while the stump-tailed and pig-tailed macaques showed a high degree of horizontal overlap. The pig-tailed macaque also segregated from the rhesus macaque, particularly at smaller spatial scales. Both these species used the forest interior almost exclusively for their daily activities but whenever they came into direct contact, as, for instance, when both fed on the fruits of *Castanopsis indica* in winter, they generally avoided confrontation. The pig-tailed and stump-tailed macaques similarly overlapped extensively in horizontal space but avoided aggressive encounters when they encountered one another. Comparable patterns of spatial overlap, without overt conflict, have been reported between sympatric bonnet and lion-tailed macaques in the Western Ghats (Singh & Roy, 2011) and between pig-tailed and rhesus macaques in Bangladesh (Feeroz, 2012) as well.

Supporting evidence for this qualitative note from our current work is provided, and partly complicated, by our quantitative home-range analysis. The rhesus and stump-tailed macaques’ home ranges were essentially separate, consistent with the contrasting habitat tolerances of a disturbance-adaptable edge species and a forest-interior specialist. Horizontal overlaps between the pig-tailed and rhesus macaques were also low, despite these two species having the most similar diets. The highest home-range overlap was, instead, recorded between the pig-tailed and stump-tailed macaques.

The overlap between the home ranges of these two forest species, however, was strongly asymmetric rather than mutual: it was driven largely by one of the stump-tailed macaque group’s much larger home range—roughly six times that of either pig-tailed macaque groups—within which the two latter ranges were largely nested, rather than by any close correspondence in habitat preference between the two species. As a small range nested within a much larger one will appear to overlap heavily regardless of whether the two species actually select similar habitats, the presence of an extensive horizontal overlap alone cannot be taken as evidence of niche similarity between the two concerned species. This is why it was important to examine the positions of pig-tailed and stump-tailed macaques on the other comparative axes of vertical stratum use, dietary profiles or temporal behavioural activity patterns to discern whether the two species were, in fact, effectively partitioning resources despite sharing so much horizontal space. The rhesus macaque’s association with the forest edge and surrounding matrix habitat, meanwhile, more plausibly reflected a comparatively higher tolerance of habitat disturbance than active avoidance of the other two species, an alternative explanation with its own implications for managing edge- and matrix habitats around the sanctuary.

Where horizontal space was shared, the three macaques diverged in their use of the vertical forest strata. Pig-tailed macaques, for example, used the highest canopy layers, more than did the other two species, to feed on the fruits of large trees such as *Ficus* spp., *Castanopsis indica*, and *Garcinia sopsopia*, and to forage for insects in the middle canopy, dominated by *Vatica lanceaefolia*; Feeroz (2012) reported similar canopy use by pig-tailed macaques in Bangladesh. The stump-tailed macaque, in contrast, was predominantly terrestrial, consistent with its principal food plants, including certain shrubs and young bamboo shoots, being low-growing while the rhesus macaque used an intermediate height range: almost exclusively arboreal inside the forest to possibly avoid predators, such as leopards and pythons, but largely terrestrial when foraging and travelling in the edge or matrix habitats. Although insects made up a substantial part of the diet of both pig-tailed and rhesus macaques, the two species foraged for them at different heights, middle canopy versus lower canopy respectively. This vertical stratification, well documented in other primate and vertebrate communities (Emmons, 1980; Estrada & Coates-Estrada, 1985; Ungar, 1996; Singh et al., 2000; Sushma & Singh, 2006; Singh & Roy, 2011; Feeroz, 2012), together with the horizontal space overlap patterns discussed above, showed that spatial partitioning operated along both axes examined in this study, consistent with findings from other sympatric congeneric primates (Cords, 1986; Sushma & Singh, 2006; Rakotondranary & Ganzhorn, 2011; Feeroz, 2012).

### Foraging and Feeding Differences Between the Three Macaque Species

All the three study macaques devoted considerable time to foraging, feeding, and moving in search of food resources, which were consistently patchily distributed in the sanctuary but which could have otherwise brought the study groups into frequent interspecies competition over these resources. Differences in the diet composition and food niche breadth of the two taxa, however, appear to have reduced this potential for competition. Rhesus macaques, generalists able to exploit a wide variety of forest- and matrix resources, displayed the broadest food niche breadth of the three species and consistently so across seasons. Pig-tailed macaques, on the other hand, had a narrow, but seasonally stable, niche breadth, relying on a select set of resources in both seasons. The stump-tailed macaque’s niche breadth was narrowest of all in the dry season, when it fed almost exclusively on two readily available resources: *Forrestia (= Amischotolype) mollissima* and *Schizostachyum polymorphum* (see also Stotrabhashyam et al., 2023); surprisingly, this pattern was the opposite of that reported for other nonhuman primates, where diet breadth typically widens when food is scarce (Schoener, 1971), before broadening in the wet season to include a greater diversity of food plants.

The three species also differed in the specific food resources and the localities they visited in search of these resources. In the dry season, for example, the stump-tailed macaque relied on the aforementioned shrub *Forrestia mollissima* and the bamboo *Schizostachyum polymorphum* in the forest interior while the rhesus macaque fed on insects, and on *Acacia auriculiformis* and *Lantana camara* in the surrounding vegetation matrix and tea plantations; the two species also roosted in different trees, stump-tailed macaques mainly in interior-forest *Ficus* spp., *Artocarpus chama*, and *Dipterocarpus retusus*, and rhesus macaques in bamboo species and *Lagerstroemia flos-reginae* near the forest edges or human habitations. In the wet season, both species used the fruits of *Sapium baccatum* and *Litsea monopetala* but at different localities within their home ranges: the stump-tailed macaques in the forest interior and degraded sanctuary areas, the rhesus macaques at the sanctuary edge and in the surrounding matrix.

Food niche overlap was highest between the rhesus and pig-tailed macaques in both seasons and especially so in the dry season, when both fed heavily on insects and on the abundant, long-fruiting *Castanopsis indica*. Despite this dietary overlap and their extensive overlap in horizontal space, discussed above, the two species achieved niche partitioning through markedly different use of vertical space, a further illustration of how a high overlap on one niche axis can coincide with strong differentiation on another, as has been observed elsewhere despite similar high overlaps in the use of abundant resources (Terborgh, 1984; Sushma & Singh, 2006).

### Time-Activity Budgets and Temporal Behavioural Differentiation

The study groups of the three congeneric species in Hollongapar differed significantly in how they distributed the six examined behavioural activities—foraging, feeding, resting, moving, and social interactions—across the day. Notably, these differences persisted after controlling for season and adult sex class, accounting for variation among groups and the non- independence of observations within days, and excluding the sparsely sampled 0500–0700 h observational period. As temporal differentiation of behavioural performance is a measure of when behavioural activities occur rather than simply how time is allocated to these behaviours overall, the significant species × time-block interactions that our study reveals provide more direct evidence of behavioural differentiation than the annual behavioural activity percentages alone. Stump-tailed macaques exhibited the highest probabilities of foraging-related movement from mid-morning onwards, consistent with observed travel between feeding sites or dispersed resources although between-group variation in foraging and feeding allocation was appreciable: the group-level variance component was greater than zero for feeding and social interactions, with feeding accounting for 41.9% of observational time in one stump-tailed macaque group but only 25.5% in the other, also cautioning us against treating the species-level budget as fixed. Rhesus macaques rested more than the other species through most of the day, which may reflect efficient exploitation of concentrated resources or an energy-conservation strategy rather than lower ecological activity, consistent with their known behavioural flexibility (Jaman & Huffman, 2013). Pig-tailed macaques devoted the largest share of observations to foraging and feeding, particularly in the morning, potentially reflecting food items that are more dispersed, difficult to locate or that require prolonged handling. The three species also differed in when social interactions peaked across the day, likely reflecting species-specific patterns of resting, group cohesion, and affiliation, although the early observational period estimates should be interpreted cautiously, given the low sample sizes at those times.

Together, these patterns suggest that time-activity budgets and temporal behavioural differentiation reflect a combination of food distribution, dietary requirements, habitat structure, and social demands that can differ even among closely related, sympatric species (van Schaik et al., 1983; Menon & Poirier, 1996). Such temporal differences in behavioural strategies may reduce how often the three congeneric species require comparable level of resources simultaneously over the day and could complement the spatial and dietary separation that have emerged in our study, consistent with findings from other sympatric nonhuman primate communities (Sushma & Singh, 2006; Zhou et al., 2014; Howell et al., 2026). Differences in such temporal behavioural activity allocation alone, however, do not by themselves demonstrate ecological niche partitioning, as species could use similar resources at different time periods or show similar time-activity budgets while differing in the food species chosen or the forest strata or actual locations used; we, therefore, treat the present results as evidence of temporal behavioural differentiation, which could be partially representative of a suite of extant coexistence mechanisms and which could complement the partitioning of space use and dietary profiles, reported above, rather than as independent proof of partitioning by themselves.

### Caveats and Limitations of Our Study

A few limitations should be considered when interpreting the results we have discussed in this paper.

1. Each macaque species was represented by two groups alone; group identity entered the vertical-stratum and time-activity budget models as a random effect. The variance components having been estimated from six groups, however, the observed species differences could partly reflect group-specific influences.
2. None of the study individuals of any of the macaque species groups could be reliably identified and all the species were largely sampled on non-overlapping days; short-term environmental variation could thus be confounded with species identity and residual temporal autocorrelations persist in the study.
3. The habitat spatial overlap estimates were method-sensitive: those macaque pairs, whose home ranges were separate, when analysed by the minimum convex polygon method, exhibited up to ∼12% overlap under 95% kernel density estimation, indicating low-intensity peripheral overlaps rather than absolute exclusion of their home ranges.
4. Finally, we did not examine intra-specific competition, which intuitively needs to exceed inter-specific competition levels for stable coexistence (Chesson, 2000) and our long-term explorations could not distinguish competitive exclusion (Miller & Max, 2025) from stochastic extinction as a cause for the local extinction of the Assamese macaque in Hollongapar (Sharma et al., 2012b). We must, therefore, admit that the ecological niche partitioning, described here, could potentially represent a transient state of current ecological being at our disturbed study site.

## Conclusions

The results from this investigation provide strong evidence that niche partitioning along spatial, dietary, and temporal axes may have helped the three studies congeneric macaque species coexist in this small, resource-limited fragment of the Upper Brahmaputra Valley, and may have helped the nonhuman primates populations to persist here through more than a century of isolation. Only a few key food resources appear to be used extensively in this forest patch by the more ecologically specialised species, with the pig-tailed macaques primarily depending on *Ficus* spp., *Castanopsis indica*, and *Garcinia sopsopia*, and the stump-tailed macaques on *Forrestia* (=*Amischotolype*) *mollissima* and *Schizostachyum polymorphum*. These resources, and the structural complexity of the forest that sustains them, thus become critical to the continued survival of both macaque species in Hollongapar and in similar fragments as well; maintaining and, where necessary, restoring these plant taxa should, therefore, be a conservation priority in the coming years. The more generalist rhesus macaque, in contrast, relies extensively on the vegetation matrix surrounding the forest, and managing this matrix could both improve the survival prospects of this species and, at the same time, reduce its interactions with the more specialised species and with the people in the surrounding villages, who often complain about the depredations caused by this macaque in their homes and fields..

More broadly, Hollongapar is one of only a handful of sites where three congeneric macaques persist in sympatry (Sharma et al., 2012b; Zhou et al., 2026), and our findings suggest that fine-scale partitioning across multiple ecological niche axes, rather than any single axis alone, is what allows such an assemblage to persist in a fragment, with an increasingly degrading area of less than 21 km², consistent with the view that coexisting species must differ sufficiently across their ecological requirements to persist together over time (MacArthur & Levins, 1967). As tropical forests worldwide are drastically reducing to small, isolated patches of fragmented habitats (Haddad et al., 2015; McKinney, 2025), understanding how closely related species share the diminishing resources of these shrinking habitats will be central to predicting which species can coexist in what remains and which, like the Assamese macaques of Hollongapar, will first die out in our last forests of the Anthropocene.

## Acknowledgements

We express our sincere gratitude to our late field collaborators, Dilip Boruah and Noren Bhuyan for their unstinting support in the field and to M D Madhusudan for his warm friendship and guidance over the years. We express our sincere gratitude to the Assam Forest Department for granting us permission to conduct this study in the Hollongapar Gibbon Sanctuary.

## Author Contributions

Narayan Sharma: Conceptualisation, Methodology, Formal Analysis, Investigation, Data Curation, Writing – Original Draft, Writing- Review and Editing, Visualisation, Funding Acquisition. Anindya Sinha: Conceptualisation, Methodology, Resources, Writing – Review and Editing, Supervision, Project Administration, Funding Acquisition.

## Data Availability Statement

The data used to support the findings of this study are available from the corresponding author upon reasonable request.

## Declarations

### Ethical Statement

All results reported in this article adhered to the legal research requirements of India, with our observational field research being conducted with permission from the Chief Wildlife Warden of the Assam State Forest Department, Guwahati, India. Our research did not require the handling of any nonhuman individuals, dead or alive, and adhered to the principles for the ethical treatment of primates, in accordance with the IPS Code of Best Practices for Field Primatology.

## Supplementary Materials

**Supplementary Table S1.** GLMM results for vertical height use among the three sympatric macaque species. Generalised linear mixed models (Tweedie family, log link) have been fit with species as a fixed effect and group identity as a random intercept, for each of three height datasets: Overall height, Feeding height, Foraging height. The significance of species differences has been assessed via likelihood-ratio tests against a random-effects-only null model. The estimated marginal means (EMMs) have been back-transformed from the log scale. The pairwise contrasts have used Bonferroni-adjusted *p*-values, with three comparisons per dataset.

| Dataset | Number of Scans (N) | Df<sup>*</sup> | χ<sup>2†</sup> | ***p***<sup>‡</sup> | Troop ID Random-Effect SD<sup>¶</sup> |
|---|---|---|---|---|---|
| **Overall height** | 33,989 | 2 | 14.58 | **< 0.001** | 0.171 |
| **Feeding height** | 10,444 | 2 | 17.48 | **< 0.001** | 0.104 |
| **Foraging height** | 969 | 2 | 12.17 | **0.002** | 0.534 |
| Dataset | Pig-tailed Macaque EMM<sup>*</sup> (95% CI<sup>†</sup>) | Rhesus Macaque EMM (95% CI) | Stump-tailed Macaque EMM (95% CI) |  |  |
| **Overall height** | 15.68 (12.35–19.90) | 7.16 (5.64–9.09) | 4.06 (3.20–5.15) |  |  |
| **Feeding height** | 16.83 (14.52–19.51) | 8.47 (7.29–9.83) | 5.68 (4.90–6.59) |  |  |
| **Foraging height** | 11.56 (5.49–24.34) | 4.97 (2.30–10.73) | 0.27 (0.10–0.74) |  |  |
| Dataset | Pairwise Contrast | Ratio | z<sup>*</sup> | *p* (Bonferroni-adjusted)<sup>†</sup> |  |
| ***Overall height*** |  |  |  |  |  |
|  | Pig-tailed macaque versus Rhesus macaque | 2.19 | 4.55 | **< 0.001** |  |
|  | Pig-tailed macaque versus Stump-tailed macaque | 3.86 | 7.85 | **< 0.001** |  |
|  | Rhesus macaque versus Stump-tailed macaque | 1.77 | 3.30 | **0.003** |  |
| ***Feeding height*** |  |  |  |  |  |
|  | Pig-tailed macaque versus Rhesus macaque | 1.99 | 6.40 | **< 0.001** |  |
|  | Pig-tailed macaque versus Stump-tailed macaque | 2.96 | 10.16 | **< 0.001** |  |
|  | Rhesus macaque versus Stump-tailed macaque | 1.49 | 3.71 | **0.0006** |  |
| ***Foraging height*** |  |  |  |  |  |
|  | Pig-tailed macaque versus Rhesus macaque | 2.33 | 1.55 | 0.367 |  |
|  | Pig-tailed macaque versus Stump-tailed macaque | 42.86 | 5.86 | **< 0.001** |  |
|  | Rhesus macaque versus Stump-tailed macaque | 18.42 | 4.53 | **< 0.001** |  |

<sup>*</sup>Df = Degrees of freedom; <sup>†</sup>χ² = Chi-square estimate; <sup>‡</sup>*p*-values in bold indicate significance at p < 0.05; <sup>¶</sup>SD = Standard deviation.
<sup>*</sup>EMM = Estimated marginal means; <sup>†</sup>95% CI = 95% confidence intervals.
<sup>*</sup>z = Wald z statistic; <sup>†</sup>*p*-values in bold indicate contrasts, significant at α = 0.05 after Bonferroni correction.
The pig-tailed macaque versus rhesus macaque contrast for Foraging height (*p* = 0.367) was not significant, in contrast to the corresponding scan-level Dunn's test, reflecting the reduced effective sample size (two groups per species), after group-level non-independence was accounted for.

**Supplementary Table S2.** Time-activity budgets of adult individuals of the three macaque species at Hollongapar Gibbon Sanctuary, given as the percentage of scan records, assigned to each broad activity category, for each species and for each study group.

| **Species** | **Group** | **n<sup>\*</sup>** | **Foraging** | **Feeding** | **Resting** | **Moving** | **Social Interactions** | **Other Behaviour** |
|---|---|---|---|---|---|---|---|---|
| **Pig-tailed macaque** | All groups | 6,471 | 6.1 | 37.3 | 14.5 | 33.9 | 5.4 | 2.8 |
|  | PTM1 | 4,394 | 6.7 | 34.6 | 15.4 | 34.1 | 6.4 | 2.8 |
|  | PTM2 | 2,077 | 4.9 | 42.9 | 12.5 | 33.6 | 3.4 | 2.7 |
| **Rhesus macaque** | All groups | 11,061 | 0.9 | 27.8 | 29.0 | 31.6 | 8.1 | 2.6 |
|  | RM1 | 6,394 | 0.8 | 30.9 | 29.6 | 29.2 | 7.6 | 1.9 |
|  | RM2 | 4,667 | 1.0 | 23.6 | 28.1 | 35.0 | 8.6 | 3.6 |
| **Stump-tailed macaque** | All groups | 16,717 | 2.9 | 30.2 | 10.3 | 48.9 | 5.7 | 2.0 |
|  | STM1 | 11,925 | 3.5 | 25.5 | 11.3 | 51.1 | 6.0 | 2.5 |
|  | STM2 | 4,792 | 1.3 | 41.9 | 7.8 | 43.5 | 4.8 | 0.6 |

<sup>\*</sup>n = Total number of scans.
Group codes follow Table 1. Percentages are of all classified adult scan records and sum to 100% across the six behavioural categories. The n values are marginally lower than the scan totals in Table 2 because seven records lacked one of the six behavioural categories and were excluded.

**Supplementary Table S3.** Random-effects variance components from the beta-binomial mixed models of temporal behavioural activity allocation, fitted with group identity and group-date identity as nested random intercepts.

| **Model set** | **Behaviour** | **Group SD<sup>\*</sup>** | **Group-day SD** | **LR<sup>†</sup>** | ***p*<sup>‡</sup>** |
|---|---|---|---|---|---|
| **Full model** | Foraging | 0.243 | 0.845 | 1.66 | 0.099 |
|  | Feeding | 0.201 | 0.614 | 3.98 | **0.023** |
|  | Resting | 0.0002 | 0.450 | 0.00 | 0.500 |
|  | Moving | 0.073 | 0.449 | 0.33 | 0.284 |
|  | Social Interactions | 0.272 | 0.780 | 3.17 | **0.037** |
| **Sensitivity model** | Foraging | 0.246 | 0.838 | 1.78 | 0.091 |
|  | Feeding | 0.206 | 0.614 | 4.34 | **0.019** |
|  | Resting | 0.0002 | 0.461 | 0.00 | 0.500 |
|  | Moving | 0.080 | 0.458 | 0.44 | 0.254 |
|  | Social Interactions | 0.267 | 0.779 | 3.01 | **0.041** |

<sup>\*</sup>SD = Standard deviation; <sup>†</sup>LR = Likelihood-ratio test statistic; <sup>‡</sup>*p*-values in bold indicate significance at p < 0.05
SD values are on the logit scale. Here, LR and *p* test whether group-level variance exceeds zero, by comparison with a model, omitting the group-level intercept; as this test lies on a parameter boundary, *p* is reported as a 50:50 mixture of chi-squared distributions.
The full models included six time blocks from 05:00 h to 17:00 h; the sensitivity models excluded the sparsely sampled 05:00–07:00 h period. All ten models converged, with convergence code of 0 and positive-definite Hessian. The group-level term for Resting was estimated at effectively zero, with SD = 0.0002, indicating a singular fit for that variance component; the corresponding models are, therefore, equivalent to those omitting the component.

